# Highly parallel genome variant engineering with CRISPR/Cas9 in eukaryotic cells

**DOI:** 10.1101/147637

**Authors:** Meru J. Sadhu, Joshua S. Bloom, Laura Day, Jake J. Siegel, Sriram Kosuri, Leonid Kruglyak

## Abstract

Direct measurement of functional effects of DNA sequence variants throughout a genome is a major challenge. We developed a method that uses CRISPR/Cas9 to engineer many specific variants of interest in parallel in the budding yeast *Saccharomyces cerevisiae*, and to screen them for functional effects. We used the method to examine the functional consequences of premature termination codons (PTCs) at different locations within all annotated essential genes in yeast. We found that most PTCs were highly deleterious unless they occurred close to the C-terminal end and did not interrupt an annotated protein domain. Surprisingly, we discovered that some putatively essential genes are dispensable, while others have large dispensable regions. This approach can be used to profile the effects of large classes of variants in a high-throughput manner.

## Main text

Understanding the functional effects of DNA sequence variants is of critical importance for studies of basic biology, evolution, and medical genetics, but measuring these effects in a high-throughput manner is a major challenge. One promising avenue is precise editing with the CRISPR/Cas9 system, which allows generation of DNA double-strand breaks (DSBs) at genomic sites matching the targeting sequence of a guide RNA (gRNA). Recent studies have used CRISPR libraries to generate many frameshift mutations genome-wide through faulty repair of CRISPR-directed breaks by nonhomologous end-joining (NHEJ) (*1*). We sought to adapt this approach to precise variant engineering. Generation of precise gene edits by CRISPR/Cas9 requires introduction of a repair template that can be used to direct repair through homology-directed repair (HDR) pathways (*2*), in the process incorporating the desired sequence variants present on the template into the genomic locus. Generating many uniquely edited cells in parallel thus requires each cell to receive the correct gRNA-repair template pair. We devised an approach that accomplishes such pairing by encoding gRNA targeting sequences and their corresponding repair templates in *cis* on oligonucleotides generated in bulk with high-throughput synthesis. These oligonucleotide libraries are then used to generate pools of plasmids pairing the two components for delivery into yeast cells. A similar method was recently reported in bacteria (*3*). We used this approach to understand the consequences of one important class of genetic variants: premature termination codons (PTCs).

PTCs interrupt the open reading frames (ORFs) of protein-coding genes. Such mutations are generally expected to have strong deleterious effects, either by abrogating or changing the functions of the encoded proteins or by causing mRNA degradation through the nonsense-mediated decay (NMD) surveillance pathway. More than 10% of annotated pathogenic human variants are PTCs (*4*, *5*). Nonetheless, our understanding of the detrimental effects of PTCs is incomplete, particularly when they occur near the 3′ ends of genes. Such mutations may not shorten the encoded proteins sufficiently to affect their function, and often escape NMD.

We first tested gene editing that employs a plasmid-encoded paired gRNA and repair template (figure 1a) by targeting eight specific PTCs to the *S. cerevisiae* genome. *S. cerevisiae* has a naturally high propensity to repair DSBs through HDR (*6*), which we enhanced by using a yeast strain in which NHEJ is abolished by a deletion of the *NEJ1* gene (*7*) (Supplementary Table 1). For each targeted mutation, we sequenced the corresponding genomic locus in thousands of transformed yeast cells. In all eight cases, the desired mutation was present in >95% of sequencing reads, demonstrating the high efficiency of this strategy (Table 1).

**Figure 1:**
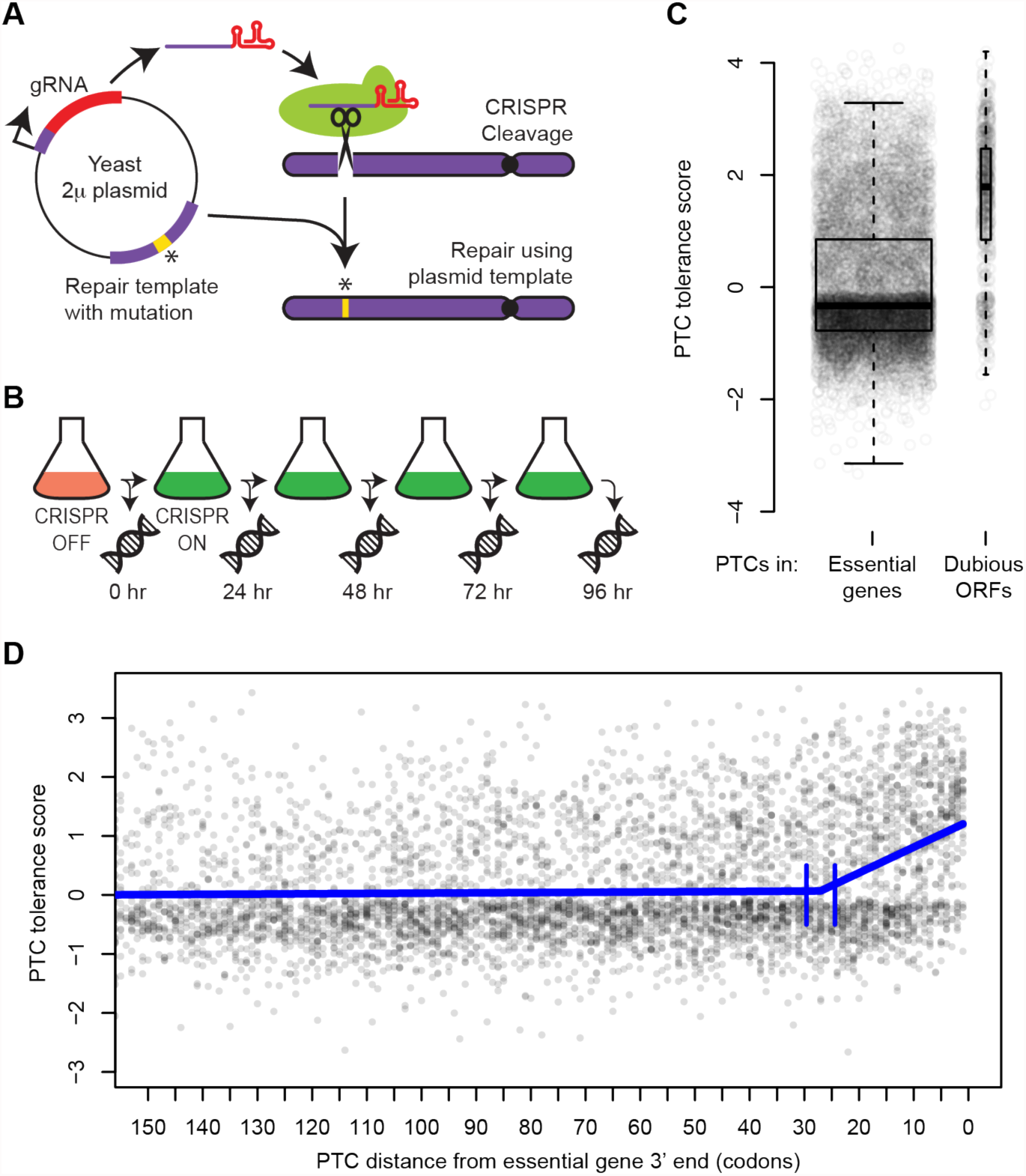
Measuring the effects of engineered PTCs in essential genes. (A) Schematic of pairing of CRISPR gRNA and repair template on plasmids. (B) Experimental design. Following Cas9 induction, DNA was extracted every 24 hours. At each time point, edit-directing plasmids were quantified by sequencing. (C) Tolerance score for each tested PTC in essential genes and dubious ORFs, with overlaid boxplots. P < 2 x 10^-16,^ Wilcoxon rank test. (D) Scatterplot of PTC tolerance scores versus distance in codons from the 3′ ends of essential genes. The thick blue line shows a segmented regression fit. Vertical blue lines indicate the 95% confidence interval for the boundary between the segments.

**Table 1:** Assessing the efficiency of edit-directing plasmids. Outcomes of directed mutations at eight loci in *nej1Δ* cells, as determined by paired-end Illumina reads of PCRs of genomic DNA at each locus.

|  | Expected edit | Unedited | Mismatch | Indel |
| --- | --- | --- | --- | --- |
| *ho*-G582Stop | 98.51% | 0.08% | 1.40% | 0.00% |
| *his2*-E308Stop | 99.83% | 0.07% | 0.10% | 0.00% |
| *mnd1*-V219Stop | 99.35% | 0.53% | 0.12% | 0.00% |
| *spo11*-F381Stop | 95.56% | 4.24% | 0.19% | 0.00% |
| *spo13*-P252Stop | 99.67% | 0.20% | 0.13% | 0.00% |
| *ste3*-P469Stop | 99.75% | 0.12% | 0.13% | 0.00% |
| *can1*-G121Stop | 99.81% | 0.06% | 0.13% | 0.00% |
| *can1*-G70Stop | 99.80% | 0.03% | 0.17% | 0.00% |

We next scaled up the approach by using large-scale oligonucleotide synthesis to generate a pool of over 10,000 distinct paired gRNA-repair template plasmids (Supplementary figure 1). These plasmids targeted PTCs to different sites in 1034 yeast genes considered essential for viability (*8*, *9*). Each gene was targeted at 10 sites, chosen with a preference for sites closer to the 3′ end (Supplementary Figure 2). We transformed yeast in bulk with this plasmid pool. After inducing Cas9 expression, we collected millions of surviving transformed cells every 24 hours for four days (figure 1b). PTCs that disrupt the function of genes essential for viability are expected to drop out of the pool over time, while those that do not are expected to persist.

**Figure 2:**
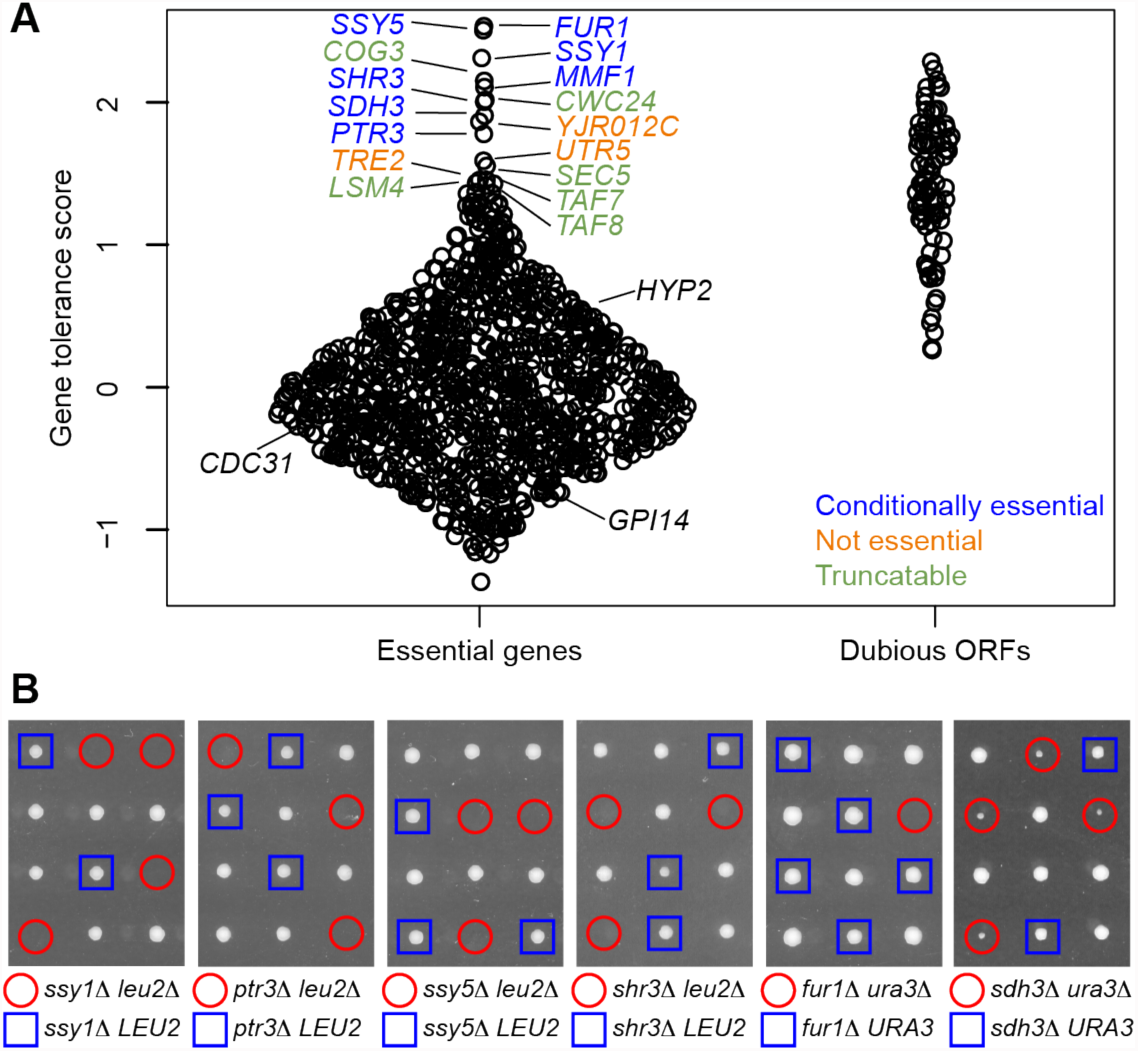
PTC tolerance of genes. (A) Gene tolerance scores for essential genes and dubious ORFs, shown as a violin plot that displays the individual data points. (B) Analysis of conditionally essential genes in yeast tetrads. Each vertical set of four colonies corresponds to the four haploid meiotic products from a diploid yeast strain. Each diploid was heterozygous for a deletion mutation of interest and for an interacting mutation. Haploid colonies carrying the deletion of interest are highlighted in red or blue based on their genotype at the interacting locus. Absence of a visible colony (first five panels) indicates a lethal interaction; small colonies (last panel) indicate an interaction causing poor growth.

We determined the abundance of each edit-directing plasmid at each time point by bulk sequencing, and computed a “PTC tolerance score” based on the persistence of each plasmid over the duration of the time-course experiment (Materials and Methods). As controls, we used a set of 90 “dubious ORFs,” which were originally annotated as genes but later reclassified due to lack of conservation and ascribable function (*10*). As expected, PTCs in essential genes were much less tolerated than those in dubious ORFs (Wilcoxon rank test, P < 2 x 10^-16)^ (figure 1c). As a further control, 71 sites in essential genes were targeted with two plasmids that had the same gRNA but different repair templates, only one of which introduced a PTC.

Plasmids that introduced a PTC were significantly less tolerated (Supplementary Figure 3) (paired t-test t = 6.5, P = 8 x 10^-9^), showing that the observed phenotypic effects are predominantly due to specific introduction of the desired mutations, rather than repair-template-independent Cas9 activities.

**Figure 3:**
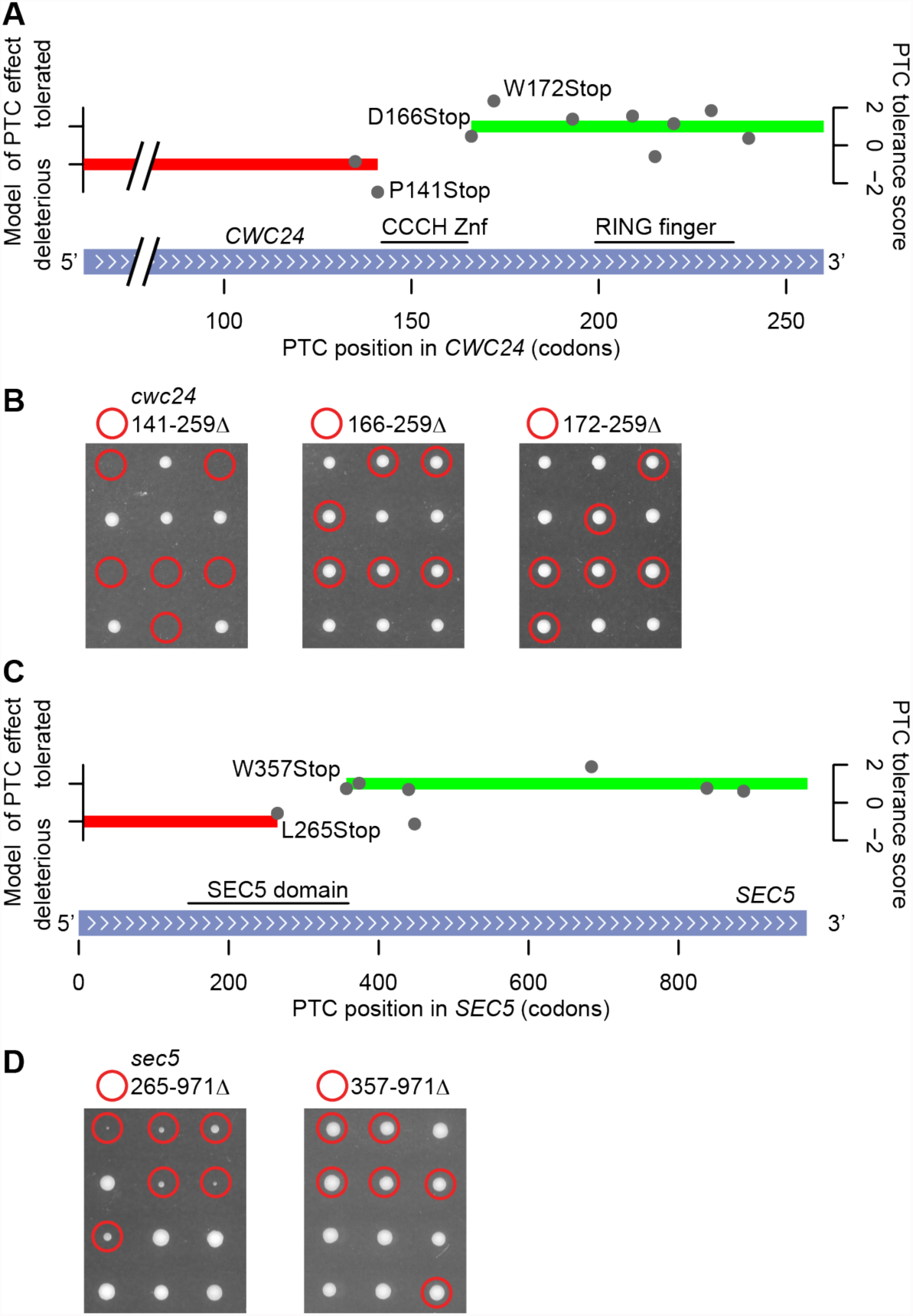
Selected truncatable essential genes. (A) Tolerance scores for 10 PTCs in *CWC24* are shown by gray circles; red and green bars indicate HMM calls of ‘deleterious’ and ‘tolerated’, respectively. The RING finger and CCCH Znf domains of Cwc24 are highlighted. (B) Analysis of deleterious and tolerated truncations of *CWC24* in yeast tetrads, displayed as in figure 2. Deletions of the last 88 and 94 codons of *CWC24* are tolerated (middle and right panels), while deletion of the last 119 codons is not (left panel). (C) Tolerance scores for eight PTCs in *SEC5* are shown by gray circles; red and green bars indicate HMM calls of ‘deleterious’ and ‘tolerated’, respectively. The Pfam-annotated “SEC5 domain” is highlighted. (D) Analysis of deleterious and tolerated truncations of *SEC5* in yeast tetrads. Deletion of the last 615 codons of *SEC5* is tolerated (right panel), while deletion of the last 707 codons is not (left panel).

One possibility for the observed PTC intolerance is that most truncations of essential genes fatally disrupt protein function. Another possibility is that NMD removes most transcripts carrying PTCs, which is fatal in the case of PTCs in essential genes. We tested these alternatives by introducing PTCs in a strain that is NMD-deficient (*11*). PTCs in this strain were similarly deleterious (Supplementary Figure 4) (χ^2^ = 1.66, P = 0.20) (Supplementary Table 8), suggesting that protein truncation, rather than degradation of transcripts via NMD, explains the observed PTC intolerance.

Although most PTCs in annotated essential genes were highly deleterious, some appeared to be tolerated. We examined the relationship between tolerance scores and locations of PTCs. PTCs were generally deleterious when located more than 27 codons away from the gene end (figure 1d). Within the 27 terminal codons, the tolerance scores rose toward the 3′ end. PTCs were also more tolerated if they did not interrupt or remove an annotated protein domain (*12*) (χ^2^ = 317.2, P = 5.86 x 10^-71^) (Supplementary figure 5, Supplementary Table 8). PTCs that disrupted protein domains tended to be deleterious even when they fell close to gene ends. Evolutionary conservation of the truncated region among related yeast species (*13*) was also significantly but more weakly predictive of PTC tolerance (χ^2^ = 49.8, P = 1.66 x 10^-12^) (Supplementary figure 5).

We built a model to more precisely delineate dispensable 3′ ends of essential genes. While our experiment is not designed to comprehensively rule out the existence of small dispensable C-termini, it is interesting to note that 517 genes did not appear to tolerate any tested PTCs (Supplementary figure 6), in some cases even very close to their ends. For instance, we confirmed that Pob3, involved in nucleosome remodeling during DNA replication and transcription, does not tolerate loss of its last two amino acids (Supplementary Figure 6), which are adjacent to the nuclear localization sequence (*14*). We also confirmed that PCNA, required for the processivity of DNA polymerase, did not tolerate the loss of its last five amino acids. This highly conserved region is part of the binding surface of PCNA used for protein-protein interactions (*15*). In contrast to the highly PTC-intolerant genes, 101 genes tolerated five or more PTCs, suggesting that these genes have large dispensable C-termini (Supplementary Figure 6). We computed the overall tolerance of PTCs for each gene and observed considerable variation among genes (figure 2a). A gene ontology enrichment analysis (*16*) showed that genes encoding proteins with catalytic activity were significantly less PTC-tolerant than other genes (Kolmogorov-Smirnov test, Bonferroni corrected P = 0.0024) (Supplementary table 10, Supplementary Figure 7), while genes with functions relating to mRNA splicing and processing were significantly more PTC-tolerant (Kolmogorov-Smirnov test, Bonferroni corrected P = 0.0017).

To better understand why some genes annotated as essential could tolerate many PTCs, we closely examined the 16 most PTC-tolerant genes (figure 2a). These genes included *SSY1, PTR3,* and *SSY5*, the three members of the SPS (Ssy1-Ptr3-Ssy5) plasma membrane amino acid sensor system (*17*), as well as *SHR3*, required for SPS cell-surface localization (*18*). Defects in SPS function compromise leucine uptake, and the strain originally used to determine which genes are essential is deficient in leucine biosynthesis and thus requires leucine uptake, which explains the lethality of SPS mutations in this strain (*19*,*20*). We confirmed that deletions of these genes were viable in yeast that could synthesize leucine, but lethal in yeast that could not (figure 2b). Similarly, the PTC-tolerant gene *FUR1* is required for the utilization of exogenous uracil (*21*), and uracil biosynthesis is also disrupted in the strain used to annotate essential genes. We confirmed that *FUR1* is only essential in yeast which cannot synthesize uracil (figure 2b), consistent with previous synthetic lethality results (*22*). Unexpectedly, we also observed poor growth of yeast with deletions of both *URA3* and the PTC-tolerant gene *SDH3* (figure 2b), a member of the mitochondrial inner membrane protein translocase complex (*23*), which suggests that proper uracil utilization may involve an unknown mitochondrial function. These examples illustrate that genes not universally essential for yeast viability can appear essential in a specific genetic background.

Another PTC-tolerant gene, *MMF1*, encodes a protein homologous to the widely conserved RidA, which processes toxic imines produced during isoleucine biosynthesis in *Salmonella* (*24*). We noticed that deletion of *MMF1* is viable under our growth conditions, but not under those previously used to define the set of essential genes (Supplementary figure 8). Thus in this case, the lethality of a gene disruption is determined by a gene-environment interaction. We hypothesize that *MMF1* is essential in yeast when isoleucine is synthesized, but not when it is taken up from the growth medium. Importantly, the isoleucine transporters Bap2 and Bap3 are poorly expressed under the growth conditions used to annotate essential genes, but more highly expressed under the growth conditions used in our experiments (*25*).

Three additional PTC-tolerant genes were misannotated as essential because their deletion disrupts the function of a nearby essential gene. *YJR012C* overlaps the 5′ end of the PTC-intolerant essential gene *GPI14* (Supplementary figure 9, figure 2a). Close examination of *YJR012C* RNA sequencing and ribosome footprinting data (*26*) indicates that the start position of *YJR012C* is misannotated (Supplementary figure 9). Deletion of *YJR012C* from its true start at M76 to the end of the ORF was viable, confirming *YJR012C* is not essential. Similarly, close examination of *UTR5* revealed that it overlaps the TATA box of the essential gene *HYP2* (*27*); deletion from the 34th codon of *UTR5* spared the TATA box and was viable (Supplementary figure 9). Finally, deletion of *TRE2* is inviable due to its effects on the neighboring essential gene *CDC31* (*28*). These examples illustrate the value of PTC introduction for characterization of gene essentiality.

Six PTC-tolerant essential genes encode proteins with large dispensable C-terminal regions. One striking case is *CWC24*, a highly conserved member of the spliceosome. Cwc24 has a CCCH-type Zinc finger domain (Znf) and a RING-type Znf domain. Analysis of the effect of PTCs in *CWC24* suggested the RING finger domain was dispensable while the CCCH Znf was essential (figure 3a), which we confirmed by engineering *CWC24* truncations (see also Wu *et al.*, 2016 (*29*)). It is interesting to note that a PTC after the RING finger domain of the essential (*30*) human homolog of *CWC24*, RNF113A, is viable (*31*). Four other PTC-tolerant genes, *TAF7*, *TAF8*, *COG3*, and *LSM4*, have been reported to tolerate large truncations (*32*–*35*). We verified that *SEC5*, a 971-amino acid member of the essential exocyst complex (*36*), tolerates truncation of at least 615 amino acids (figure 3b). Our observation that 101 genes tolerated five or more PTCs suggests that many additional genes have dispensable C-terminal regions.

Our results improve the annotation of essential genes in the well-studied yeast genome. We discovered several cases of genes that appeared to be essential as a consequence of the specific strain and growth conditions originally used to test viability of gene deletions. A deletion screen in a different yeast isolate also revealed examples of conditionally essential genes (*37*). Applying our approach and related methods (*38*) in a diverse set of isolates and growth conditions will further refine the core set of essential yeast genes.

PTCs are prioritized in studies of human genetics because of the high likelihood that they abolish gene function. Our results suggest that PTCs are most likely to be deleterious when they disrupt annotated protein domains or truncate more than 27 amino acids, and these criteria may improve filtering of candidate causal variants. We observed that NMD did not make a strong contribution to PTC tolerance. This result is consistent with recent findings that NMD in yeast acts most strongly on transcripts with PTCs toward their 5′ ends (*39*). PTCs near the ends of human genes are also likely to escape NMD according to the 50-base-pair rule (*40*) (Supplementary figure 10), and our criteria may be especially useful for predicting their effects.

In our study we carried out a pooled screen of the functional effects of approximately 10,000 directed mutations. Our method can be used to assess the functional effects of any desired nucleotide variants in a highly parallel manner. The ability to profile the impact of broad classes of alleles, including missense and regulatory variants, will enable a more fine-grained understanding of the relationship between genotypes and phenotypes.

## Acknowledgements

We thank Kruglyak laboratory members, F. Albert, M.P. Hughes, and J. Rine for helpful discussion, R. Cheung and E. Pham for technical assistance, and G. Church for plasmids. Funding was provided by the Howard Hughes Medical Institute and NIH grants R01 GM102308 (L.K.) and F32 GM116318 (M.J.S.).

